# AFG3L2-mediated proteolysis restricts mitochondrial biogenesis and gene expression in hypoxia

**DOI:** 10.1101/2024.09.27.615438

**Authors:** Srikanth Chandragiri, Nils Grotehans, Yvonne Lasarzewski, Maria Patron, Thomas MacVicar, Yohsuke Ohba, Steffen Hermans, Elena Rugarli, Hendrik Nolte, Thomas Langer

## Abstract

Mitochondria are metabolically rewired in hypoxia when cells switch to glycolytic growth. In addition to the well-established role of transcriptional and translational programs, there is increasing evidence that post-translational mechanisms contribute to the rapid adaptation of the mitochondrial proteome to hypoxia. Here, we have used a proteomic survey to define how the m-AAA protease AFG3L2, a proteolytic complex in the inner mitochondrial membrane, regulates mitochondrial proteostasis. Our experiments identify a broad spectrum of mitochondrial substrate proteins and show that AFG3L2 is activated in hypoxia along an HIF1α-mTORC1 signaling axis. AFG3L2-mediated proteolysis restricts mitochondrial biogenesis and gene expression by degrading proteins, which are involved in mitochondrial protein import, mitochondrial transcription, mRNA processing, mRNA modification and stability, and RNA granule formation. Our experiments highlight the important contribution of proteolytic rewiring of the mitochondrial proteome for the adaptation to low oxygen tension and shed new light on the pathophysiology of several neurodegenerative disorders associated with mutations in *AFG3L2*.

## Introduction

Mitochondria are central metabolic hubs that dynamically adapt their structure and function in response to changing metabolic demands and environmental cues(1, 2). Metabolic reprogramming of mitochondria occurs during development, cell differentiation, immune responses, disease, and ageing, and is associated with changes in mitochondrial content and shape (3–6). Transcriptional and translational programs promote the adaptation of mitochondria to different metabolic demands and stress conditions, such as hypoxia, where HIF family transcription factors switch cells from oxidative to glycolytic metabolism and maintain redox homeostasis (7). However, there is increasing evidence for the importance of post-translational control mechanisms in mitochondrial reprogramming downstream of transcriptional regulation(8, 9). Mitochondrial proteases are emerging as key regulators of these processes. mTORC1 inhibition in hypoxia or in starved cells leads to the activation of the AAA protease YME1L (also known as i-AAA protease) in the inner mitochondrial membrane (IMM), which broadly rewires the mitochondrial proteome to acutely limit mitochondrial biogenesis and support anaplerotic reactions, such as pyrimidine synthesis. YME1L-mediated proteolysis supports xenograft growth of pancreatic ductal adenocarcinoma cells and maintains the adult neural stem cell pool in the brain (10, 11).

The m-AAA protease is a ubiquitously expressed AAA protease in the IMM, whose subunits are homologous to YME1L but expose their catalytic sites to the matrix space (12). It forms homo-oligomeric hexameric complexes with AFG3L2 and hetero-oligomeric complexes with AFG3L2 and SPG7(13). Mutations in these subunits cause several neurodegenerative diseases, including autosomal dominant spinocerebellar ataxia, recessive hereditary spastic paraplegia, and dominant optic atrophy (DOA12) (14–16). The identification of several substrate proteins of the m-AAA protease revealed crucial functions of the m-AAA protease in mitochondria (17, 18). The protease regulates mitochondrial ribosome assembly by processing the ribosome subunit MRPL32 and OXPHOS assembly by degrading TIMMDC1, a respiratory complex I assembly factor (19, 20). Degradation of the glutathione transporter SLC25A39 by AFG3L2 maintains mitochondrial glutathione homeostasis (21, 22). In addition, the m-AAA protease degrades EMRE, an essential subunit of the mitochondrial calcium uniporter MCU, thereby controlling Ca^2+^ uptake into mitochondria, whereas degradation of the H^+^/Ca^2+^ channel TMBIM5 (GHITM) by the m-AAA protease limits Ca^2+^ efflux from the organelle (23–25).

Here, we performed comprehensive proteomic studies to elucidate how the m-AAA protease affects mitochondrial proteostasis. We identify novel proteolytic substrates of AFG3L2 and show that mTORC1-regulated proteolysis by AFG3L2 largely reshapes the mitochondrial proteome and limits mitochondrial gene expression in hypoxia. Thus, our results reveal a critical role for the m-AAA protease in mitochondrial programming and highlight the importance of proteolytic rewiring of mitochondria in response to hypoxia.

## Results

### AFG3L2 broadly rewires the mitochondrial proteome

To define how AFG3L2 affects the mitochondrial proteome, we performed data-independent (DIA) mass spectrometry (MS)-based proteomics on wildtype (WT) and AFG3L2-deficient HeLa cells. We quantified 8527 protein groups, of which 925 are part of MitoCarta 3.0 (26). Statistical analysis revealed significantly lower levels of 435 mitochondrial proteins in *AFG3L2*^-/-^ cells (Figure 1A; Supplementary Table 1). Determination of log2 fold change distributions of all detected mitochondrial proteins in WT and *AFG3L2^-/-^* cells revealed an overall reduced mitochondrial mass in cells lacking AFG3L2 (Figure 1B). In contrast, 139 mitochondrial proteins accumulated significantly in *AFG3L2^-/-^* cells (Figure 1A, C). RNA sequencing of WT and *AFG3L2^-/-^*cells showed that the transcription of 58 genes was increased in the absence of AFG3L2, which may explain their accumulation in AFG3L2-deficient cells (Figure 1D). However, the transcription of 81 genes corresponding to accumulating proteins was either not significantly altered or reduced in *AFG3L2^-/-^*cells compared to WT cells (Figure 1D), indicating that the encoded proteins represent potential substrates of AFG3L2.

**Figure 1.**
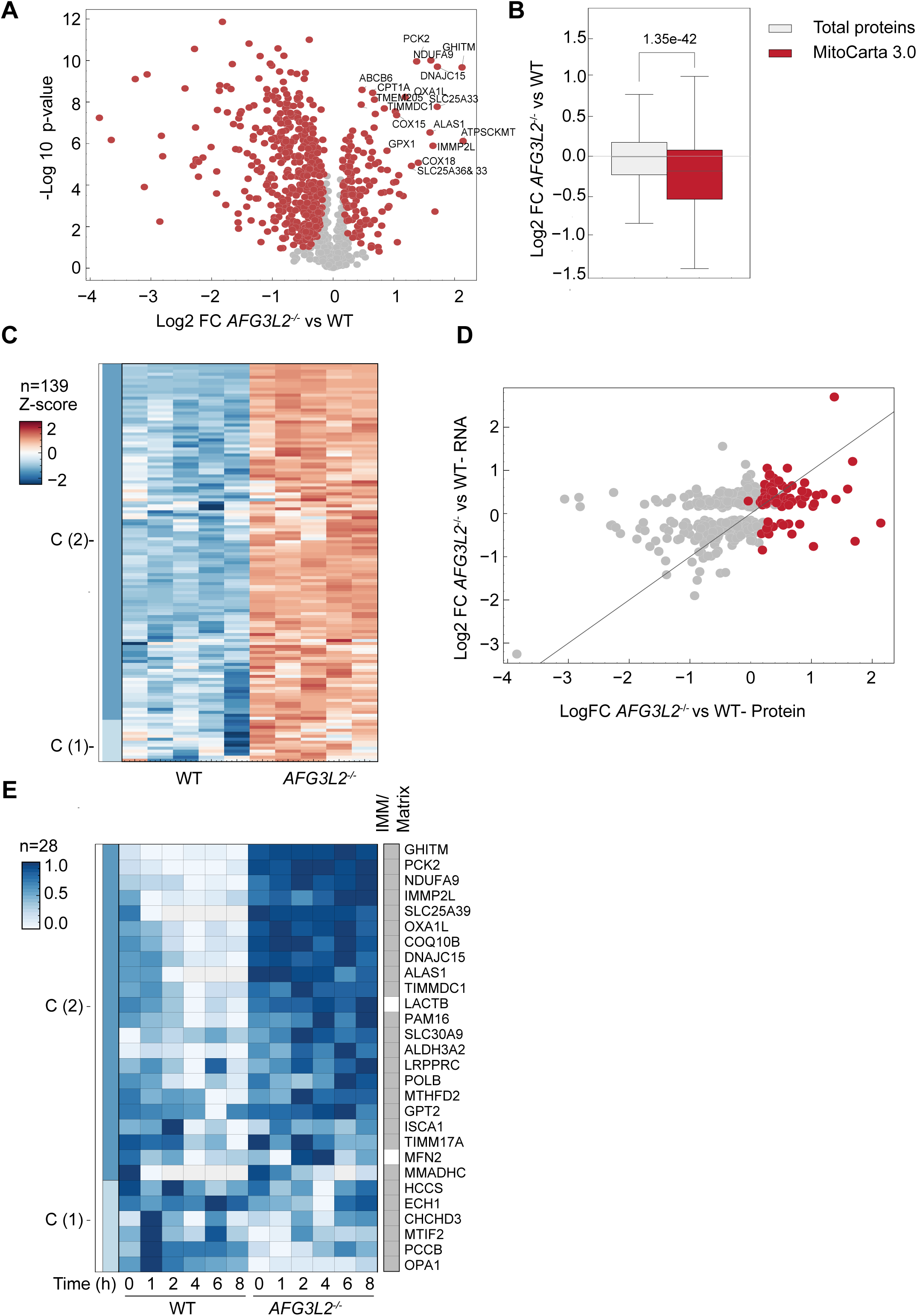
AFG3L2 broadly regulates mitochondrial proteostasis. **A.** Volcano plot depicting the log^2^ fold changes (FC) in protein abundances in *AFG3L2^-/-^* versus wildtype (WT) HeLa cells and the negative log_10_ p-value of two-tailed t-tests. Significantly changed proteins (permutation-based FDR<0.05, s0=0.1, #permutations=500) are highlighted in red. n=5 per genotype. **B.** Boxplot analysis of the cellular proteome of WT and *AFG3L2^-/-^* cells. The distribution of relative protein abundances (log_2_ FC between *AFG3L2^-/-^*and WT cells) are shown for the total proteome (grey) and for mitochondrial proteins according to MitoCarta3.0 (42). The p-value of a two-tailed t-test is indicated. **C.** Z-score heatmap of log_2_-transformed label-free quantification (LFQ) intensities of significantly increased mitochondrial (FDR<0.05) proteins in *AFG3L2^-/-^* HeLa cells. n=5 per genotype. **D.** Scatter plot showing log_2_ FC between WT and *AFG3L2^-/-^* cells in protein abundance (x-axis) and mRNA abundance (y-axis). Only proteins are only shown if the corresponding mRNA was detected and significantly changed in the RNAseq experiment. The line represents the identity curve f(x)=y. Proteins accumulating in *AFG3L2^-/-^*cells (Figure 1C) are highlighted in red. **E.** Heatmap showing the log_2_-transformed LFQ intensities of mitochondrial proteins scaling the highest intensity to 1 and the lowest intensity to zero, after aggregating the biological replicates using the median. Significantly changed proteins were identified using a mixed linear model using the time, genotype and replicate as variables on the peptide data (random effect). P-values below 0.01 for the interaction p-value (time x genotype). Only proteins whose abundance decreased with time are shown (negative slope, param time).

To exclude that translational regulation causes protein accumulation, we examined the stability of mitochondrial proteins after inhibition of cytosolic translation by cycloheximide (CHX). We determined changes in the steady-state levels of mitochondrial proteins over time after CHX addition and identified 384 proteins, whose stability differed significantly between WT and *AFG3L2^-/-^* cells using a mixed linear model (Supplementary Figure 1A). Filtering for mitochondrial proteins using MitoCarta 3.0 revealed that 28 mitochondrial proteins were stabilized in *AFG3L2^-/-^* cells (Figure 1E). The latter proteins are mainly localized in the IMM or matrix, where they are accessible to the proteolytic attack by AFG3L2, and include the previously described substrates of the m-AAA protease TMBIM5 (GHITM) (25), TIMMDC1 (20), and SLC25A39 (21, 22).

Taken together, these experiments identify multiple candidate substrates of AFG3L2 and suggest broad rewiring of the mitochondrial proteome by AFG3L2-mediated proteolysis (Supplementary Table 1). Notably, the proteomic analysis of SPG7-deficient HeLa cells lacking hetero-oligomeric m-AAA proteases did not reveal significant changes in steady-state levels of mitochondrial proteins, suggesting that homo-oligomeric AFG3L2 complexes can compensate for the loss of SPG7 in these cells (Supplementary Figure 1B).

### AFG3L2-mediated mitochondrial reprogramming in hypoxia

Oxygen deprivation induces various signaling cascades, which ultimately adapt mitochondrial metabolic functions and the defence against oxidative stress (6). Increasing evidence suggests that altered protein turnover rates allow rapid reshaping of the mitochondrial proteome in response to hypoxia, complementing regulatory circuits at transcriptional and translational levels (10, 27). Proteomic and transcriptomic analysis of cells at normoxia and hypoxia (0.5% O_2_) identified 153 mitochondrial proteins, which failed to accumulate in hypoxia despite increased transcription (Figure 2A). We have previously demonstrated that mTORC1-dependent signaling activates the *i*-AAA protease YME1L, which broadly reprograms the mitochondrial proteome to support anaplerotic reactions (10). Since we observed decreased levels of many matrix-localized proteins and additional IMM proteins in hypoxia (Supplementary Figure 2A), we reasoned that AFG3L2 may also contribute to the proteolytic rewiring of mitochondria under these conditions. Indeed, steady-state levels of 31 AFG3L2 candidate substrates were decreased in hypoxic mitochondria compared to normoxic conditions, although the corresponding mRNA levels were not reduced (Figure 2A).

**Figure 2.**
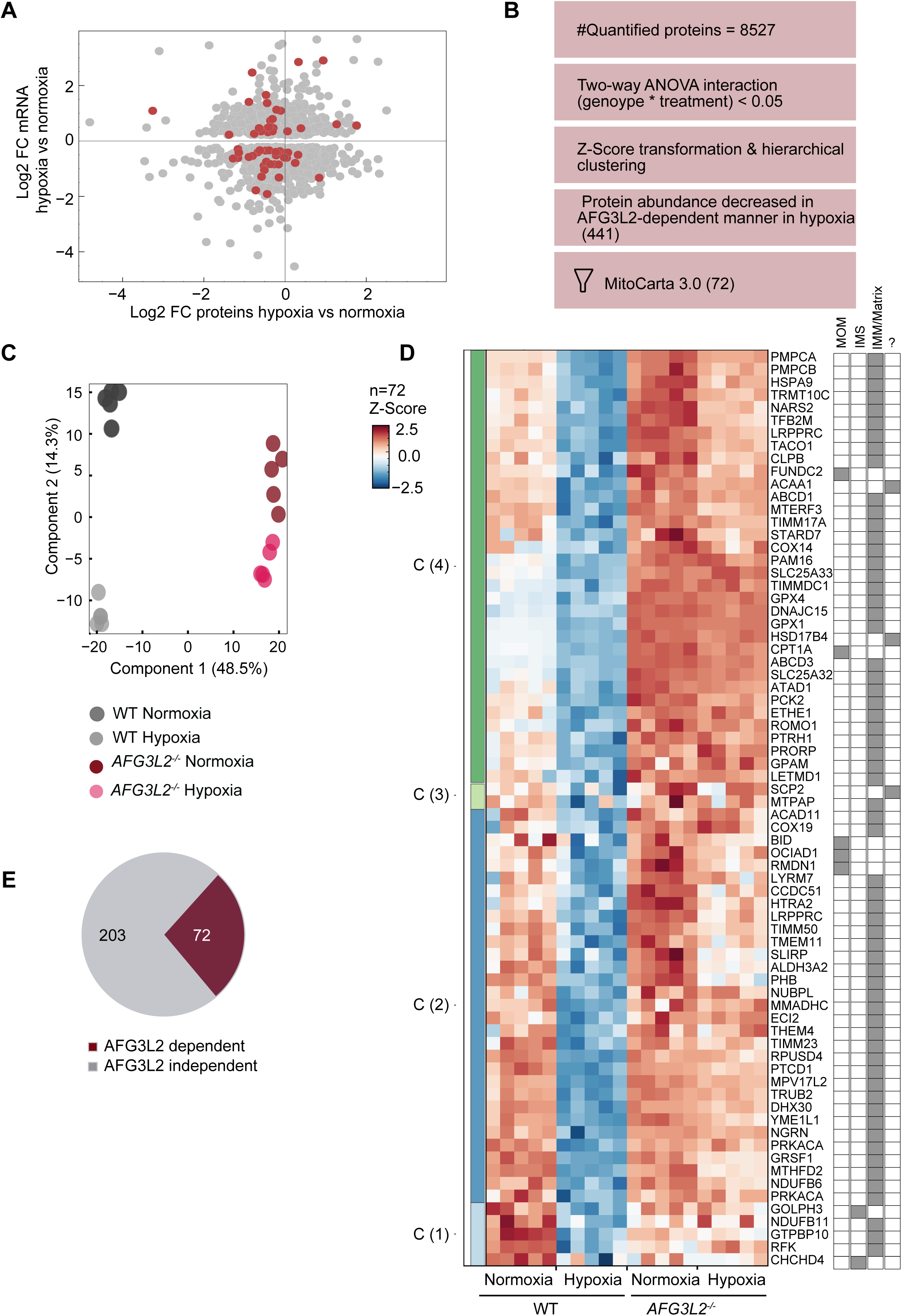
AFG3L2-mediated mitochondrial reprogramming in hypoxia. **A.** Scatter plot showing log_2_ fold changes (FC) in abundances of proteins (x-axis) or mRNAs (y-axis) in wildtype (WT) cells in hypoxia or normoxia (10). Mitochondrial proteins significantly accumulating in *AFG3L2^-/-^* cells under normoxic conditions (Figure 1C) are highlighted in red. **B.** Workflow of the proteomic analysis of WT and *AFG3L2^-/-^* cells in normoxia and hypoxia (0.5% O_2_; 16 hours). **C.** Principal component analysis (PCA) of the cellular proteome of WT and *AFG3L2*^-/-^ cells in normoxia and hypoxia. **D.** Z-score heatmap of log_2_-transformed LFQ intensities of mitochondrial proteins, whose abundance decreases significantly in WT HeLa cells in normoxia but increases or remains unchanged in *AFG3L2^-/-^* HeLa cells (n=5). Localization of proteins to mitochondria and to different mitochondrial subcompartments according to MitoCarta 3.0. **E.** Venn-diagram showing the fraction of proteins, whose level is decreased in hypoxia in an AFG3L2-dependent manner among all mitochondrial proteins with decreased abundance in hypoxic cells.

We therefore performed MS-based quantitative proteomics on WT and *AFG3L2*-deficient HeLa cells in normoxia (21% O_2_) and hypoxia (0.5% O_2_) (Figure 2B). We identified 8527 proteins, of which 925 were annotated as mitochondrial proteins according to MitoCarta 3.0 (26). Dimensionality reduction by principal component analysis (PCA) revealed segregation by genotype (component 1 48.5%) and oxygen tension (component 2 14.3%) (Figure 2C). Notably, the segregation of WT cells by component 2 was more profound than that of *AFG3L2*-deficient cells, indicating a greater proteome rewiring in response to oxygen tension in WT cells. This effect was particularly pronounced for the mitochondrial proteome, which was greatly reduced in hypoxic compared to normoxic WT cells, a response that occurred only moderately in *AFG3L2*-deficient cells (Supplementary Figure 2B).

Two-way ANOVA analysis of the proteomic dataset identified 1780 significantly regulated proteins, which were clustered into four distinct groups by hierarchical clustering using Euclidean distance (Supplementary Figure 2C). We focused on proteins, whose abundance decreased in response to hypoxia in WT cells but not in *AFG3L2*-deficient cells (Supplementary Figure 2C; cluster 1). This cluster contains 441 proteins, including 72 mitochondrial proteins, 62 of which are localized to the mitochondrial matrix or IMM (Figure 2D), including the previously described AFG3L2 substrate TIMMDC1 (20). Another known AFG3L2 substrate, the glutathione transporter SLC25A39 (28), accumulated in *AFG3L2*^-/-^ cells, but was not detected in all replicates of WT cells and therefore did not cluster with other AFG3L2 substrates (Supplementary Figure 2D). Notably, 31 proteins accumulated in *AFG3L2*^-/-^ cells under normoxic and hypoxic conditions, indicating proteolytic turnover by AFG3L2 independent of the oxygen tension (Figure 2D). Another group of 41 proteins did not accumulate in *AFG3L2*^-/-^ cells in normoxia, but was stabilized under hypoxic conditions in the absence of AFG3L2, indicating AFG3L2-dependent proteolytic degradation only in hypoxia (Supplementary Table 2).

Together, these results identify multiple candidate substrates of AFG3L2 in hypoxic mitochondria. We have observed decreased steady-state levels of 275 mitochondrial proteins in hypoxia, 72 of them (corresponding to 26%) dependent on AFG3L2 (Figure 2E), highlighting the central role of AFG3L2 for the proteolytic rewiring of mitochondria in hypoxia. It is noteworthy that we also identified 303 proteins that accumulate in hypoxia only in AFG3L2-dependent manner, including cytosolic chaperones, proteins involved in peroxisomal biogenesis, and mitochondrial proteins (Supplementary Figure 2C; cluster 3), indicating that AFG3L2 plays a pleiotropic role for cellular adaptation to low oxygen tensions.

### Posttranslational regulation of AFG3L2-mediated protein turnover

To investigate the role of the canonical hypoxic transcription factor HIF1α that is stabilized in hypoxia for the AFG3L2-dependent mitochondrial rewiring, we used the HIF1α-stabilizing compounds dimethyloxalylglycine (DMOG) and CoCl_2_ (29). Treatment with DMOG or CoCl_2_ resulted in reduced levels of the AFG3L2 substrates TIMMDC1 and SLIRP in normoxia indicating increased AFG3L2 activity (Supplementary Figure 3A). This cannot be explained by increased *AFG3L2* expression, as AFG3L2 protein levels were not altered in hypoxia (Supplementary Figure 3B). Therefore, we conclude that HIF1α regulates AFG3L2 post-translationally.

mTORC1 signaling is inhibited in hypoxia (30) and may modulate proteolysis by AFG3L2. To examine this possibility, we inhibited mTORC1 pharmacologically or by various metabolic cues and monitored the accumulation of candidate AFG3L2 substrates. Steady-state levels of AFG3L2 substrates decreased after mTORC1 inhibition with rapamycin or torin in WT but not in *AFG3L2^-/-^* cells (Figure 3A, Supplementary Figure 3C). To substantiate these findings, we used MEFs lacking TSC2, a negative regulator of mTORC1, the loss of which causes constitutive activation of mTORC1 (31). While AFG3L2 substrate proteins were reduced in WT cells under hypoxic conditions, they accumulated at similar levels under normoxia and hypoxia in *Tsc2^-/-^ cells* (Supplementary Figure 3D. Conversely, inhibition of mTORC1 by torin was sufficient to activate AFG3L2 in WT and *Tsc2^-/-^* cells (Supplementary Figure 3E).

**Figure 3.**
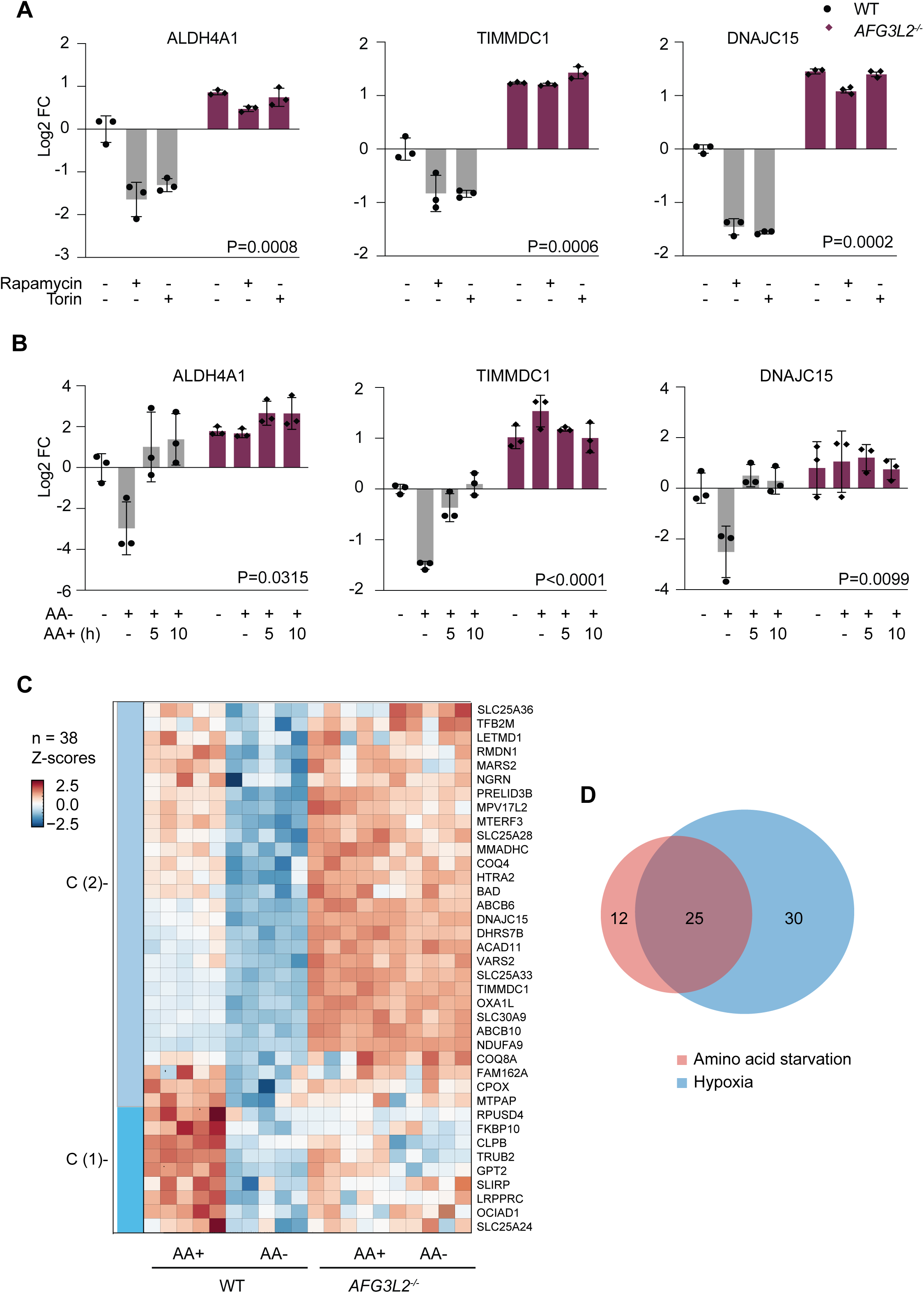
mTORC1-dependent regulation of AFG3L2. **A.** Quantification of the steady-state levels of the indicated proteins determined by immunoblotting of wildtype (WT) and *AFG3L2*^-/-^ HeLa cells treated with rapamycin (200 nM/12 hours) or torin (200 nM/12 hours) as indicated (n=3). The p-value (genotype x treatment) of two-way ANOVA analysis is indicated. See also Supplementary Figure 3C. **B.** Quantification of the steady-state levels of the indicated proteins determined by immunoblotting of WT and *AFG3L2*^-/-^ HeLa cells either starved for amino acids (-AA) or incubated with amino acids (+AA) as indicated (n=3). The p-value (genotype x treatment) of two-way ANOVA analysis is indicated. See also Supplementary Figure 3F. **C.** Z-score heatmap of log_2_-transformed LFQ intensities of AFG3L2 candidate substrate proteins in WT and *AFG3L2*^-/-^ HeLa cells, which were cultivated in the absence of amino acids for 5 hours. Proteins are shown, whose abundance significantly decreased only in cells expressing AFG3L2 (n=5). **D.** Venn-diagram showing the overlap of AFG3L2 candidate substrates identified in hypoxia (blue) or after amino acid starvation of the cells.

Similarly, we observed lower steady-state levels of selected AFG3L2 substrates in cells after amino acid starvation, which causes mTORC1 inhibition (Figure 3B, Supplementary Figure 3F). The steady-state levels of these substrates were restored in WT but not in *AFG3L2^-/-^*cells upon re-supply of amino acids (Figure 3B). Mass-spectrometric analysis of the mitochondrial proteome showed broadly increased AFG3L2-mediated proteolysis in amino acid starved cells (Figure 3C). PCA revealed segregation of the samples based on genotype and amino acid starvation, indicating that the treatment led to profound proteomic changes in both WT and *AFG3L2* cells (Supplementary Figure 3G, H). We identified 38 mitochondrial proteins whose levels were reduced in WT cells upon amino acid starvation, whereas they accumulated or were unaffected and did not respond to amino acid starvation in *AFG3L2*-deficient cells (Figure 3C). 25 of these proteins were also identified as candidate AFG3L2 substrate proteins in hypoxia (Figure 3D). Taken together, we conclude that hypoxia promotes protein degradation by AFG3L2 through a HIF1α-mTORC1-signaling axis.

### mTORC1 inhibition promotes proteolysis by AFG3L2

Since mTORC1 inhibition regulates the cytosolic translation of some mitochondrial proteins dependent on the translation initiation factor 4E-binding protein (4E-BP) (32), it is conceivable that reduced translation decreases protein levels in an AFG3L2-dependent manner. However, inhibition of mTORC1 with torin or in hypoxia decreased the steady-state levels of AFG3L2 substrates in both WT and *4e-bp^-/-^* cells, suggesting that AFG3L2 affects these proteins post-translationally (Supplementary Figures 3I, J).

Inhibition of mTORC1 induces autophagy (33), which may be affected by the loss of AFG3L2. However, the level of AFG3L2 substrates decreased upon mTORC1 inhibition in autophagy-deficient *Atg5^-/-^* MEFs, demonstrating that the regulation of AFG3L2-mediated proteolysis by mTORC1 occurs independently of autophagy (Supplementary Figures 3K, L).

To unambiguously demonstrate that mTORC1 regulates protein turnover through AFG3L2, we performed pulse SILAC (stable isotope labeling with amino acids in cell culture) experiments in WT and *Tsc2^-/-^* MEF cells (Figure 4A). Cells were labeled with N^15^-arginine and N^15^-lysine (heavy) and then transferred to media containing N^14^-arginine and N^14^-lysine (34). The synthesis of new proteins, containing light amino acids, and the degradation of pre-existing proteins leads to the decay of proteins labelled with heavy amino acids over time. We observed no systematic difference in the natural logarithmic transformation of the incorporation rates (light amino acids/(light amino acids+heavy amino acids)) between WT and *Tsc2^-/-^* cells (Supplementary Figure 4A). This indicates similar incorporation rates for the majority of proteins quantified and allows protein degradation rates to be determined by monitoring the decrease of heavy amino acids over time. Area under the curve (AUC) calculations distinguish slower (high AUC) from faster (lower AUC) protein turnover (Figure 4B).

**Figure 4.**
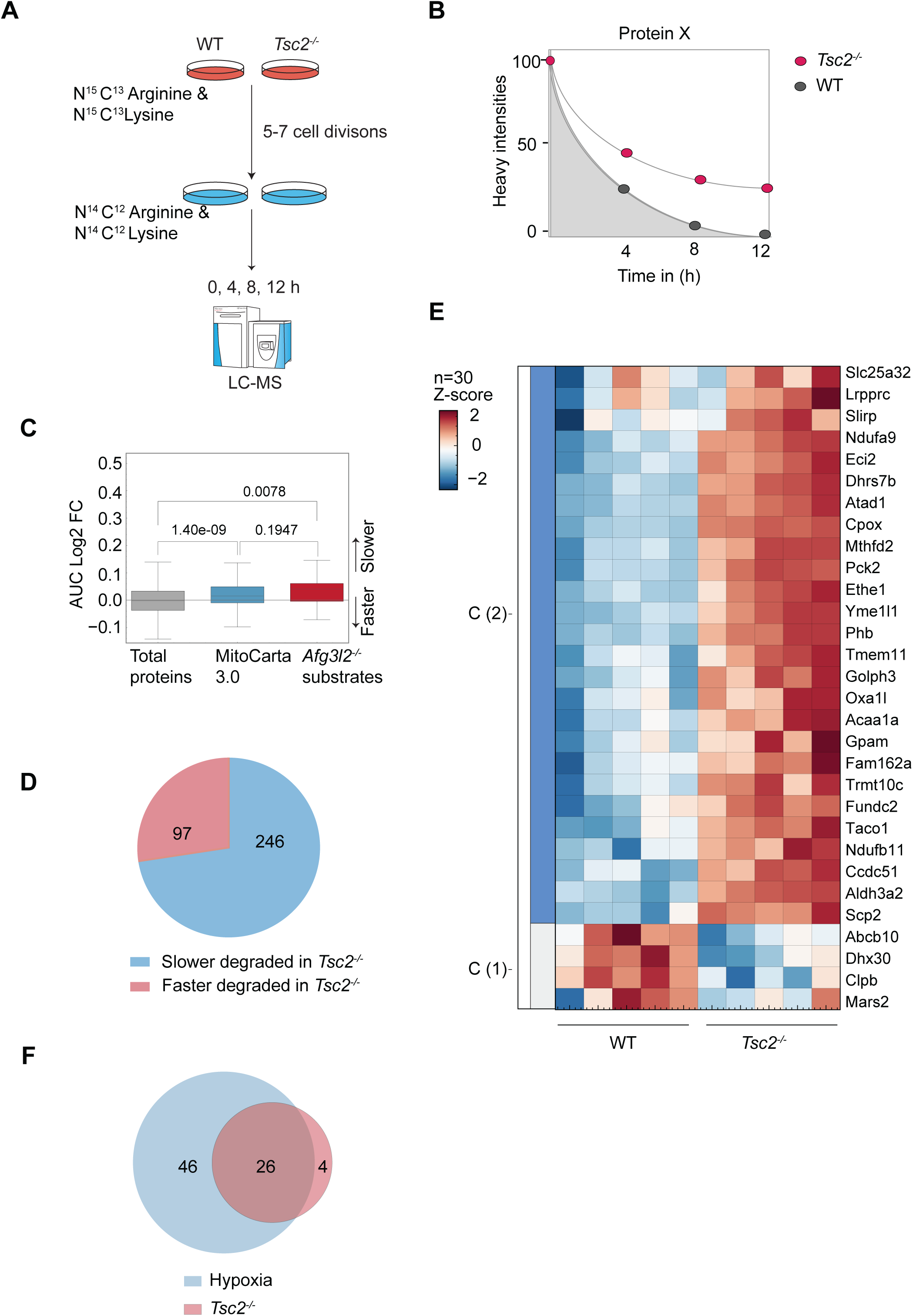
mTORC1 inhibition promotes proteolysis by AFG3L2. **A.** Workflow of the pulsed SILAC analysis. Wildtype (WT) and *Tsc2*^-/-^ MEFs were labelled with N^15^C^13^-arginine and N^15^C^13^-lysine for the indicated time and then cultivated in the presence of N^14^C^12^-arginine and N^14^C^12^-lysine. Cells were collected at the indicated times and samples were analysed by LC/MS-MS. **B.** Schematic representation illustrating the area under curve (AUC) calculation of heavy (N^15^) intensities in WT and *Tsc2^-/-^*cells. **C.** Boxplot analysis of relative protein abundances in WT and *Tsc2^-/-^*cells. The distribution of log_2_ FC in the AUC of heavy intensities of *Tsc2^-/-^* relative to WT cells. The stability of the total proteome is compared with that of the mitochondrial proteins (according to MitoCarta3.0) and the AFG3L2 candidate substrates. The p-value of a two-tailed t-test is indicated. **D.** Venn-diagram showing mitochondrial proteins with altered turnover rates in *Tsc2*^-/-^ cells and fractions of slower (blue) and faster (42) degraded proteins. **E.** Z-score heatmap of log_2_-transformed AUC heavy (N^15^) intensities of significantly changed AFG3L2 candidate substrates (Figure 2D) in *Tsc2^-/-^* cells relative to WT cells (n=5). **F.** Venn-diagram showing the fraction of AFG3L2 candidate substrates in hypoxia (blue) identified in the SILAC proteomic experiments in *Tsc2^-/-^*cells (42).

Our analysis identified 2561 significantly altered proteins that formed two clusters (Supplementary Figures 4B, C): Proteins that are stabilized upon mTORC1 activation in *Tsc2^-/-^* cells (1309 proteins; cluster 1) and proteins that were more rapidly degraded under these conditions (1251 proteins, cluster 2) (Supplementary Figure 4C). We did not observe differences in the degradation of the total proteome between WT and *Tsc2*^-/-^ cells (Supplementary Figure 4D), but the overall degradation rate of mitochondrial proteins was significantly reduced in *Tsc2^-/-^* cells (Figure 4C). Among 639 identified mitochondrial proteins in the pulse SILAC experiments, 343 proteins showed significantly altered turnover rates (Figure 4D), demonstrating the important contribution of proteolysis to the reshaping of the mitochondrial proteome by mTORC1.

To assess whether active mTORC1 inhibits AFG3L2-dependent proteolytic rewiring, we compared our list of newly identified AFG3L2 substrates in hypoxia (Figure 2D) with the subset of mitochondrial proteins that were degraded more slowly in *Tsc2^-/-^* cells (Figure 4E). Notably, 26 out of the 30 significantly altered AFG3L2 substrates identified in the pulse SILAC experiments were stabilized in *Tsc2^-/-^* cells, supporting the notion that active mTORC1 suppresses AFG3L2-dependent protein turnover (Figure 4F). Furthermore, 26 AFG3L2 substrate proteins identified in *Tsc2^-/-^* cells also accumulated in hypoxia in the absence of AFG3L2 (Figures 2D, 4F).

We conclude from these experiments that mTORC1 regulates AFG3L2-dependent proteolysis. Similar to AFG3L2, the homologous i-AAA protease YME1L is activated upon mTORC1 inhibition, which decreases phosphatidylethanolamine (PE) levels in the IMM (10, 27). However, a MS-based analysis of the mitochondrial proteome in cells lacking either the phosphatidylserine (24)-specific lipid transfer protein PRELID3B or the PS decarboxylase PISD in the IMM revealed that, in contrast to proteolysis by YME1L, decreased mitochondrial PE levels are not sufficient to promote AFG3L2-mediated protein degradation (Supplementary Figures 4E, F), suggesting a different regulatory mechanism.

### AFG3L2-dependent proteolytic rewiring limits mitochondrial transcript accumulation in hypoxia

To define the functional consequences of AFG3L2-mediated proteolysis in hypoxia, we performed a biological pathway analysis of the 72 proteins, whose steady-state levels were decreased in hypoxia in an AFG3L2-dependent manner (Figure 2D). This analysis revealed that AFG3L2 affected mitochondrial gene expression (21 out of 72 proteins), mitochondrial protein import (9 out of 72 proteins), quality control (8 out of 72 proteins), and metabolic pathways (9 out of 72 proteins) in hypoxia (Figure 5A), suggesting a pleiotropic role of AFG3L2 in mitochondria. Proteins involved in mitochondrial RNA metabolism, but not proteins associated with mtDNA maintenance or mitochondrial translation, accumulated in AFG3L2-deficient cells in hypoxia (Figure 5B). These proteins are part of mitochondrial RNA granules (MRGs) (35) and regulate mitochondrial transcription, the processing of polycistronic mRNAs as well as mRNA modification and degradation (Figure 5C, D) (36). We therefore hypothesized that AFG3L2 activation in hypoxia promotes the degradation of mitochondrial RNA metabolizing enzymes to adapt mitochondrial gene expression to low oxygen tension.

**Figure 5.**
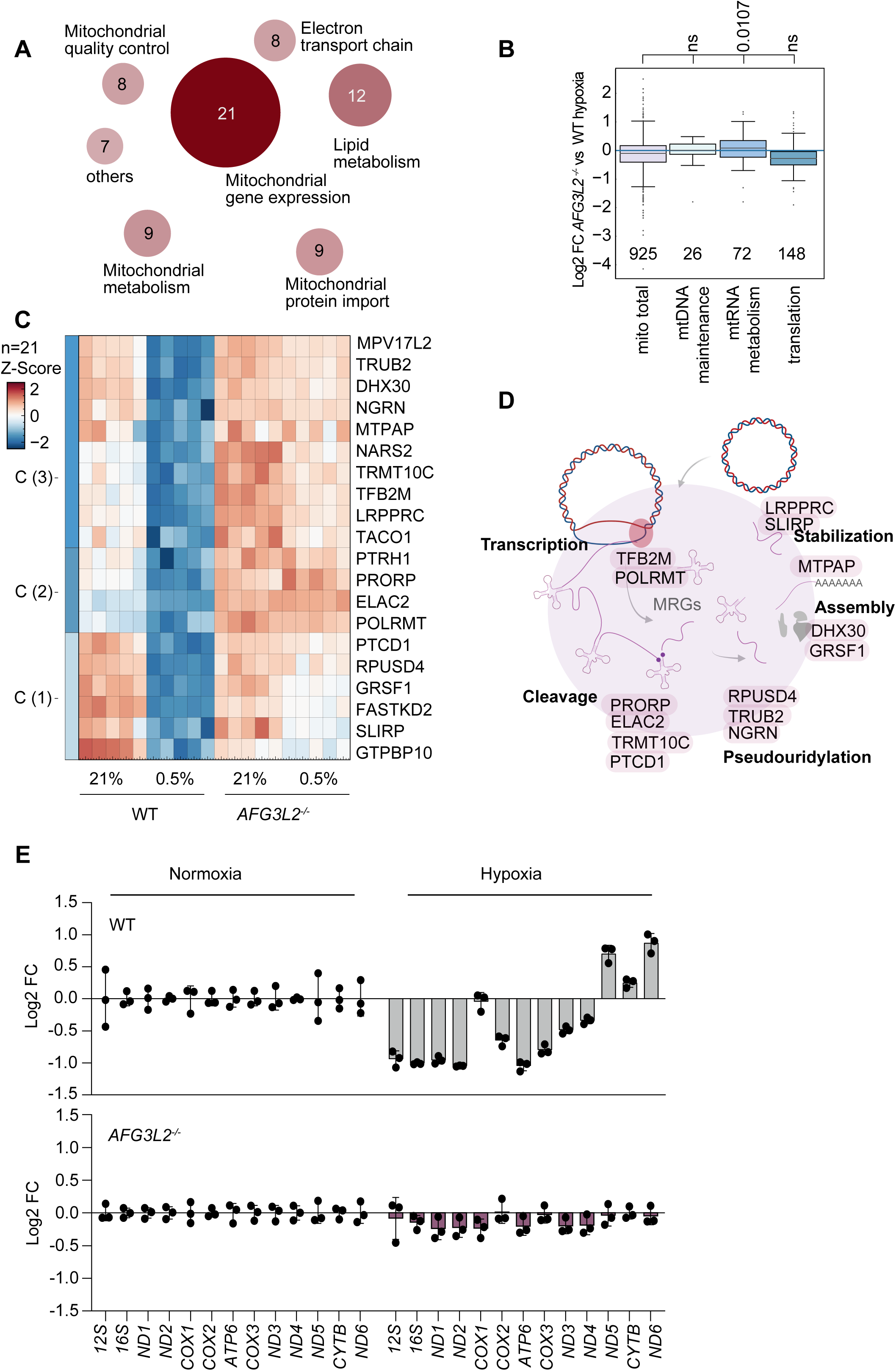
AFG3L2-dependent proteolysis limits mitochondrial gene expression in hypoxia. **A.** Pathway analysis for AFG3L2 candidate substrates in hypoxia (see Figure 2D). Circle size and color intensity indicate the representation of the pathway among AFG3L2 substrate proteins. **B.** Boxplot analysis of the mitochondrial proteome and different mitochondrial pathways in wildtype (WT) and *AFG3L2^-/-^* cells in hypoxia. The distribution of relative protein abundances (log_2_ FC between *AFG3L2^-/-^* and WT cells) are shown. The p-value of a one-way ANOVA is indicated **C.** Z-score heatmap of log_2_-transformed LFQ intensities of AFG3L2 substrate proteins involved mt-DNA gene expression in hypoxia. **D.** Schematic representation showing the role of AFG3L2 substrates in hypoxia in mitochondrial gene expression. **E.** Steady-state levels of mitochondrial transcripts in WT and *AFG3L2*^-/-^ analyzed by qRT-PCR. Log2-transformed FC in WT and *AFG3L2*^-/-^ cells relative to WT cells in normoxia and hypoxia are shown. (n=3).

Therefore, we monitored the accumulation of mitochondrial transcripts in WT and *AFG3L2^-/-^* cells under normoxic and hypoxic conditions using quantitative real-time PCR (qPCR) (Figure 5E). We observed significantly reduced RNA levels for 9 out of the 13 mitochondrial transcripts in hypoxic mitochondria when compared to normoxia (Figure 5E), suggesting that mitochondrial transcription or mRNA stability is responsive to oxygen availability and is repressed at low oxygen tension. In contrast, transcript abundance remained unchanged in hypoxic *AFG3L2*^-/-^ cells. We further confirmed that the differences in mtRNA abundance between WT and *AFG3L2*^-/-^ cells are not caused by differences in mtDNA levels in these cells, which are similar under normoxic or hypoxic conditions (Supplementary Figure 5A). Furthermore, transcripts of the known HIF1α-targets HK2 and GAPDH accumulated in both WT and AFG3L2-deficient cells under hypoxia, demonstrating that AFG3L2 is specifically required for the accumulation of mitochondrial transcripts (Supplementary Figure 5B). We therefore conclude that the proteolysis of AFG3L2 substrates associated with mtRNA metabolism limits mitochondrial transcript accumulation in hypoxia.

## Discussion

Our results highlight the critical role of mitochondrial proteolysis in the adaptation of the mitochondrial proteome to hypoxia or starvation. We observed an altered turnover rate for about half of the detected mitochondrial proteins in hypoxia. These proteins are localized in different mitochondrial compartments, suggesting the involvement of different proteases in their turnover. While we have previously shown that hypoxia activates the i-AAA protease YME1L (10), we demonstrate here that HIF1α activation and mTORC1 inhibition leads to increased proteolysis by the m-AAA protease AFG3L2. The combined action of both proteases accounts for the reduced steady-state levels of nearly 40% of the 275 reduced mitochondrial proteins in the hypoxic proteome. Other mitochondrial proteases may further increase the fraction of the mitochondrial proteome which is proteolytically rewired in hypoxia. The protease LONP1 degrades COX4-1 facilitating the adaptation of cytochrome c oxidase activity to low oxygen tension and, together with matrix CLPP, may affect the turnover of additional proteins in hypoxic cells (37).

Proteolysis allows the acute regulation of mitochondrial functions in response to hypoxia, when cells switch from oxidative to glycolytic growth with lower demands on mitochondrial OXPHOS activity (38). Proteins involved in mitochondrial gene expression are enriched among the proteolytic substrates of AFG3L2 and are acutely affected at low oxygen tension. We observed an AFG3L2-dependent decrease of mitochondrial mRNA levels in hypoxic cells, consistent with the cells switching to glycolytic growth. Our proteomic analysis identified several proteins associated with the mitochondrial RNA metabolism as substrates of AFG3L2, including LRPPRC, SLIRP and MTPAP, which regulate mRNA stability (34, 39). Recent kinetic analyses indeed suggest a critical role for proteins involved in mRNA turnover control for mitochondrial transcript accumulation and the regulation of OXPHOS (40–42). Moreover, AFG3L2 similarly affects the steady-state levels of POLRMT and TFB2M, which mediate mitochondrial transcription (43, 44), of several enzymes involved in RNA modification and the processing of polycistronic mRNAs (45, 46), and of DHX30 and GRSF1, which are required for RNA granule formation (47, 48). AFG3L2-mediated proteolysis thus restricts mitochondrial gene expression in hypoxia by interfering with mitochondrial transcription and broadly affecting the mitochondrial RNA metabolism. In addition to its compelling effect on mitochondrial gene expression, the m-AAA protease also limits the biogenesis of OXPHOS complexes post-translationally by degrading several OXPHOS assembly factors (20, 25).

We also identified several proteins involved in mitochondrial protein import as high-confidence AFG3L2 substrates, including the molecular chaperones PAM16 and DNAJC15 or the TIM23 translocase subunit TIMM17A (49). AFG3L2 activation therefore acutely limits mitochondrial protein biogenesis in hypoxia, when the cellular OXPHOS demand is reduced, which also explains the overall reduced mitochondrial mass in *AFG3L2*^-/-^ cells. It should be noted that by affecting protein biogenesis, the loss of AFG3L2 may also indirectly affect the accumulation of mitochondrial proteins and trigger protein turnover by other peptidases, such as LONP1 or CLPP, if the import of subunits of oligomeric protein complexes is reduced.

While the m-AAA protease restricts mitochondrial gene expression and OXPHOS biogenesis at low oxygen tension, it is also required for mitochondrial ribosome assembly in normoxia (19, 50). The pleiotropic functions of the m-AAA protease likely require a fine-tuned control of its proteolytic activity, which is currently poorly understood. Regulation appears to be substrate-specific, since the steady-state levels of some AFG3L2 substrates remained unchanged in hypoxia. We have previously identified the Ca^2+^/H^+^ exchanger TMBIM5 (GHITM) as an inhibitor and proteolytic substrate of AFG3L2 (and confirmed this in the present study) (25) but did not observe an increased turnover of TMBIM5 in hypoxia, suggesting other regulatory mechanisms. Similar to the i-AAA protease YME1L, inhibition of mTORC1 is sufficient to activate AFG3L2 post-translationally, but, unlike YME1L, mTORC1-dependent changes in the levels of phosphatidylethanolamine are not sufficient to promote AFG3L2-mediated proteolysis (10). We observed reduced levels of prohibitin in hypoxic mitochondria, a subunit of the PHB membrane scaffold complex, which assembles with the m-AAA protease and regulates its proteolytic activity in a substrate-specific manner (51). For instance, recruitment of the PHB complex by C9ORF12, which is associated with amyotrophic lateral sclerosis and frontotemporal dementia (FTD), was reported to inhibit AFG3L2-mediated degradation of TIMMDC1(20). It is therefore conceivable that disruption of the PHB complex in hypoxia may activate AFG3L2-mediated proteolysis. However, further studies are required to elucidate the mechanisms that lead to the increased turnover of selected substrates by the m-AAA protease in hypoxia.

Regardless, given the lower oxygen tension in the brain (52), AFG3L2 activation in hypoxia to limit mitochondrial gene expression and OXPHOS activity sheds new light on the pathophysiology of neurodegenerative disorders, which are associated with mutations in AFG3L2, and may contribute to the vulnerability of neurons to loss of AFG3L2 function.

## Material and methods

### Cell culture

HeLa and MEF cells were maintained in Dulbecco’s Modified Eagle’s Medium (DMEM) GlutaMAX (Gibco) with 10% foetal bovine serum (SIGMA) and 1 mM sodium pyruvate (Gibco) at 37°C in the presence of 5% CO_2_. All cultured cells were routinely tested for *Mycoplasma* infections. Cell growth and survival was routinely monitored by trypan blue exclusion and cell counting using the Countess automated cell counter (Thermo). Hypoxia experiments were performed in the H35 Hypoxystation (Don Whitley). Where indicated, the following compounds were added to the medium: rapamycin (200 nM), torin1 (200 nM), and cycloheximide (100 μg/ml). Amino acid starvation media were custom made and formulated according to the Gibco recipe for high-glucose DMEM, specifically omittting the amino acids. Amino acids were added concentrations according to DMEM Glutamax.

### Transfection and RNA interference

RNA interference experiments were conducted through reverse transfection of 2.5x10^5^ cells with 4 µg of esiRNA, employing Lipofectamine RNAiMax (Life Technologies). For sgRNA transfection, 2x10^5^ cells were utilized and transfected with a sgRNA plasmid (2.5 µg) using Lipofectamine LTX. Cells were harvested 48 hours after transfection.

### Generation of knockout cell lines

*Knockout* cells (*AFG3L2*^-/-^, *SPG7*^-/-^) were generated using CRISPR-Cas9 mediated gene editing in HeLa cells. Cas9 and sgRNA were expressed using the px459 vector. Individual clones were selected using immunoblotting and genomic sequencing. The following sgRNAs were used for targeting *AFG3L2* (5’-GCTGCTACCACACGCTCTTC-3’) and SPG7 (5’-GGTACATCAAGGCTAGCCGC-3’).

### Cell lysis and SDS-PAGE

Cells were harvested with ice cold phosphate buffered saline (PBS) and lysed in RIPA buffer (50 mM Tris-HCl pH7.4, 150 mM NaCl, 1% Triton X100, 0.5% DOC, 0.1 % SDS and 1 mM EDTA) for 30 min on ice. Lysates were centrifuged at 20,000x*g* for 10 min at 4°C. Protein concentrations in the supernatant fraction were determined by Bradford assay (Bio-Rad). Protein lysates were mixed with 4x lithium dodecyl sulfate (LDS) buffer (Invitrogen) and analysed by 12% SDS-PAGE and immunoblotting.

### Quantitative PCR

Genomic DNA was isolated from cell pellets using the Blood and Tissue DNA extraction kit (Qiagen). RNA was isolated from cell pellets using the RNA extraction kit (Macherey-Nagel) and 1-2 µg of RNA was reverse transcribed into cDNA using GoScript (Promega).

For measurements of mitochondrial DNA (mtDNA), 10 ng of genomic DNA were amplified using the TaqMan PCR master mix (Thermo Fisher Scientific). MtDNA levels were assessed by the delta delta ct method using *MT-RNR1*, *MT-ND4* and *MT-ND6* as mitochondrial probes and *ACTB* as nuclear DNA control. For the measurements of nuclear and mitochondrial transcripts,10 ng of cDNA was amplified using the TaqMan PCR master mix. Expression levels were calculated by the delta delta ct method, for which *B2M* was used as control.

### Proteomics sample preparation and data analysis

#### Data acquisition and analysis for CHX chase

LC-MS/MS instrumentation consisted out of an Easy-LC 1200 (Thermo Fisher Scientific) coupled via a nano-electrospray ionization source to an QExactive Hf-x mass spectrometer (Thermo Fisher Scientific) For peptide separation, an in-house packed column (inner diameter: 75 µm, length: 40 cm) was used. A binary buffer system (A: 0.1 % formic acid and B: 0.1 % formic acid in 80% acetonitrile) was applied as follows: Linear increase of buffer B from 4% to 27% within 69 min, followed by a linear increase to 45% within 5 min. The buffer B content was further ramped to 65 % within 5 min and then to 95 % within 6 min. 95 % buffer B was kept for further 10 min to wash the column. Prior each sample, the column was washed using 5 µL buffer A and the sample was loaded using 8 µL buffer A.

Full MS Spectra were acquired using a 60,000 resolution and a m/z range of 350-1650 m/z.

For MS/MS independent spectra acquisition, 48 windows were acquired at an isolation m/z range of 15 Th and the isolation windows overlapped by 1 Th. The fixed first mass was 200 m/z. The isolation center range covered a mass range of 350–1065 m/z. Fragmentation spectra were acquired at a resolution of 15,000 at 200 m/z.

The maximum injection time was set to 22ms and stepped normalized collision energies (NCE) of 24, 27, 30. The AGC target was set to 3e6 to ensure maximum filling times of the Orbitrap.

The acquired raw spectra were analyzed using the DIA-NN software (1.8) using the library FASTA-file-based analysis and the ‘robust LC’ quantification at an FDR of < 0.01. As a FASTA file, we used the Uniprot human reference proteome.

For further analysis of the data, the precursor file output from DIA-NN was utilized and the intensities were log2 transformed. The precursors were grouped to peptides using the median aggregation on the stripped peptide sequences. Then, we applied a random linear mixed model using the time and genotype as fixed effects and the peptides as random effects using the *mixedlm* function of the Python statsmodel package. The replicates were allowed to start at a different intercept. This analysis to analyse the degradation rate (slope) as well as the difference between genotypes. The interaction p-value (Genotype * Time) was utilized to identify significantly different degradation rates between genotypes. The data can be found in Supplementary file 2.

#### Data acquisition and analysis using PISD-depleted and Prelid3b^-/-^ cells

The mass spectra were acquired with the same instrumentation as described above (CHX chase). The acquired raw spectra were analyzed using the DIA-NN software (1.8.) using the library FASTA-file-based analysis and the ‘robust LC’ quantification at an pr. FDR of < 0.01. Statistical significance was accessed using a two-tailed unpaired t-test followed by a FDR correction using a permutation-based approach (5%) using the log2 transformed LFQ intensities. The data can be found in Supplementary File 5.

#### Data acquisition and analysis in hypoxia and after amino acid starvation

LC-MS/MS instrumentation consisted out of an Easy-LC 1200 (Thermo Fisher Scientific) coupled via a nano-electrospray ionization source to an Exploris 480 mass spectrometer (Thermo Fisher Scientific). The peptide separation was performed as described above.

For MS spectra acquisition, the RF Lens amplitude was set to 55%, the capillary temperature was 275°C and the polarity was set to positive. MS1 profile spectra were acquired using a resolution of 120,000 (at 200 m/z) at a mass range 320-1150 m/z and an AGC target of 1 × 106. For MS/MS independent spectra acquisition, 48 windows were acquired at an isolation m/z range of 15 Th and the isolation windows overlapped by 1 Th. The fixed first mass was to 200. The isolation center range covered a mass range of 350–1065 m/z. Fragmentation spectra were acquired at a resolution of 15,000 at 200 m/z using a maximal injection time of 22 ms and stepped normalized collision energies (NCE) of 26, 28, 30. The default charge state was set to 3. The AGC target was set to 900%. MS2 spectra were acquired as centroid spectra.

For the analysis of DIA (Data independent acquisition), we utilized DIA-NN version 1.8 (53) The library-free approach was used based on the human uniport reference proteome which predicts MS2 spectra using the neuronal network. The deep-learning option was enabled. The quantification strategy was set to ‘robust LC (high accuracy)’. The precursor range was adjusted to 330 – 1200 m/z. The RT profiling option was enabled. Otherwise, default settings were used. A two-sided t-test was applied to identify significantly different proteins as well as 2Way ANOVA using genotype (WT and AFG3L2^-/-^) and treatment (hypoxia – normoxia), (amino acids -amino acid starvation). The FDR was controlled to 5% using a permutation-based approach in the Perseus software (54).The data can be found in Supplementary File 1 and 3.

#### Data acquisition and analysis for pulse SILAC proteomics

MEF cells were grown for 5-7 doubling times in the presence of heavy amino acids (N^15^ C^13^-arginine and -lysine) and then switched to media containing light amino acids (N^14^ C^12^-arginine and -lysine). Cells were collected at different time points (0, 4, 8 and 12 hours), washed with PBS buffer and snap frozen with liquid nitrogen. Reagents used in SILAC labelling include DMEM without arginine, lysine and glutamine (Silantes, # 280001300) -100x PSG), dialyzed FBS (Gibco, # 26400044), arginine (0, 6, 10) (labeled amino acids from Silantes), and lysine (0, 4, 8) (labeled amino acids from Silantes).

Proteins were lysed and digested as described above. The raw spectra were analyzed using the Spectronaut software. The heavy and light intensities were extracted on the precursor level, log2 transformed and aggregated using the median to peptides by their stripped sequence. Then data were normalized, aligning the median of all samples (*Tsc2*^-/-^ and wild type) at each time point. The data can be found in Supplementary File 4.

### Statistics and reproducibility

All independent experiments and biological samples are represented in the graphs. Analyses were not blinded because the experiments were both conducted and analyzed by the same investigator. For proteomics measurements, the investigators were blinded to allocation during experiments, and samples were randomized. To compare two groups and treatments, a significance level of p<0.05 was considered, and a two-way ANOVA was performed. For two groups, an unpaired t-test was conducted. No statistical method was used to pre-determine the sample size. All data analyses were performed using Graph Pad Prism 10 and Instant Clue 0.12.1.

## Acknowledgements

We thank D. Diehl for excellent technical assistance. We thank MPI AGE bioinformatic facility for technical and bioinformatic assistance and MPI AGE FACS Imaging facility for technical support. NG received support from the Cologne Graduate School of Ageing Research, YO received support from a fellowship of the Japan Society for the Promotion of Science (JSPS) for research abroad, The Osamu Hayaishi Memorial Scholarship for Study Abroad and grants from the Uehara Memorial Foundation. YL was supported by the European Union’s Horizon 2020 research and innovation programme under the Marie Sklodowska-Curie grant agreement No 721757. We are grateful for the following gifts: *Tsc2^-/-^* MEFs from C. Demetriades and D. Kwiatkowski, *Atg5*^-/-^ MEFs from K. Winklhofer, WT (S/S) and 4E-BP DKO MEFs from H. McBride.

## Supplementary figure legends

**Supplementary table 1.**
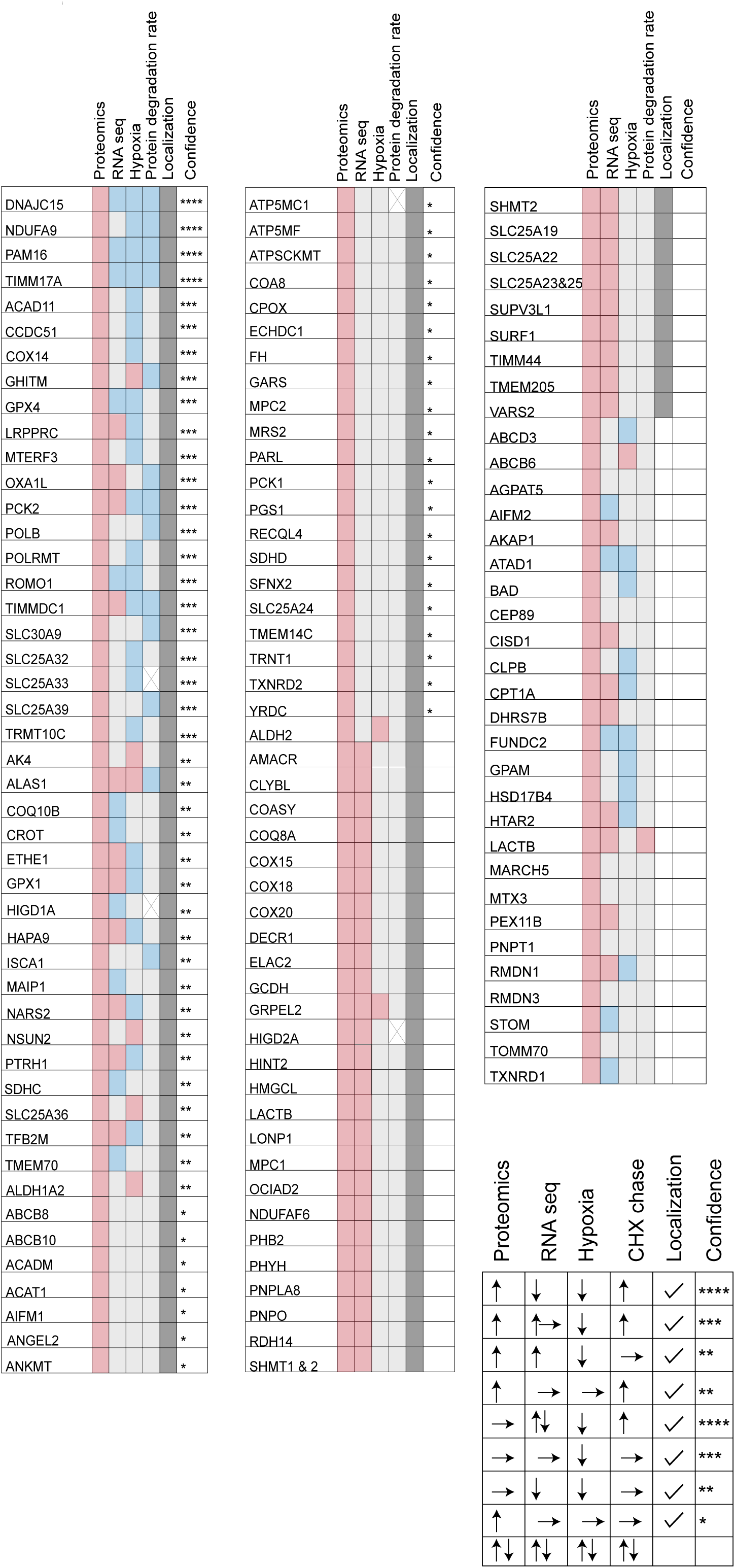
High-confidence AFG3L2 candidate substrates. Mitochondral proteins (according to MitoCarta 3.0) significantly accumulating in *AFG3L2*-/-HeLa cells (WT vs *AFG3L2*-/-) are compared with datasets of significantly altered transcript levels (RNAseq; WT vs *AFG3L2*-/-), of AFG3L2 candidate substrates in hypoxia (Figure 2D) and of protein degradation rates (Figure 1E). Upregulated proteins and increased transcript levels are indicated in red, downregulated proteins and decreased transcript levels are indicated in blue. Lower protein degradation rates are indicated in blue. The number of asterisks indicate confidence levels.

**Supplementary table 2.**
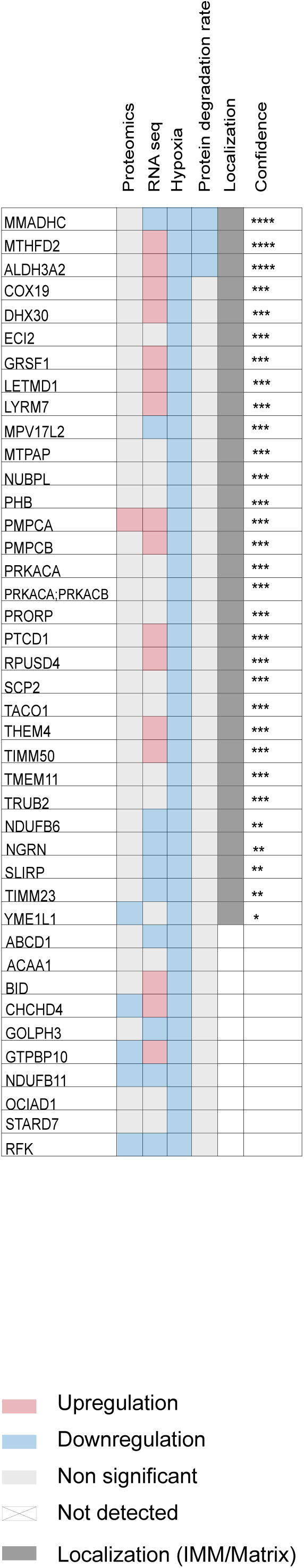
Hypoxia-specific AFG3L2 substrates whose steady-state level was not altered in *AFG3L2*-/-cells but reduced AFG3L2-dependent manner in hypoxia. The analysis was performed as described for Supplementary table 1.

**Supplementary Figure 1.**
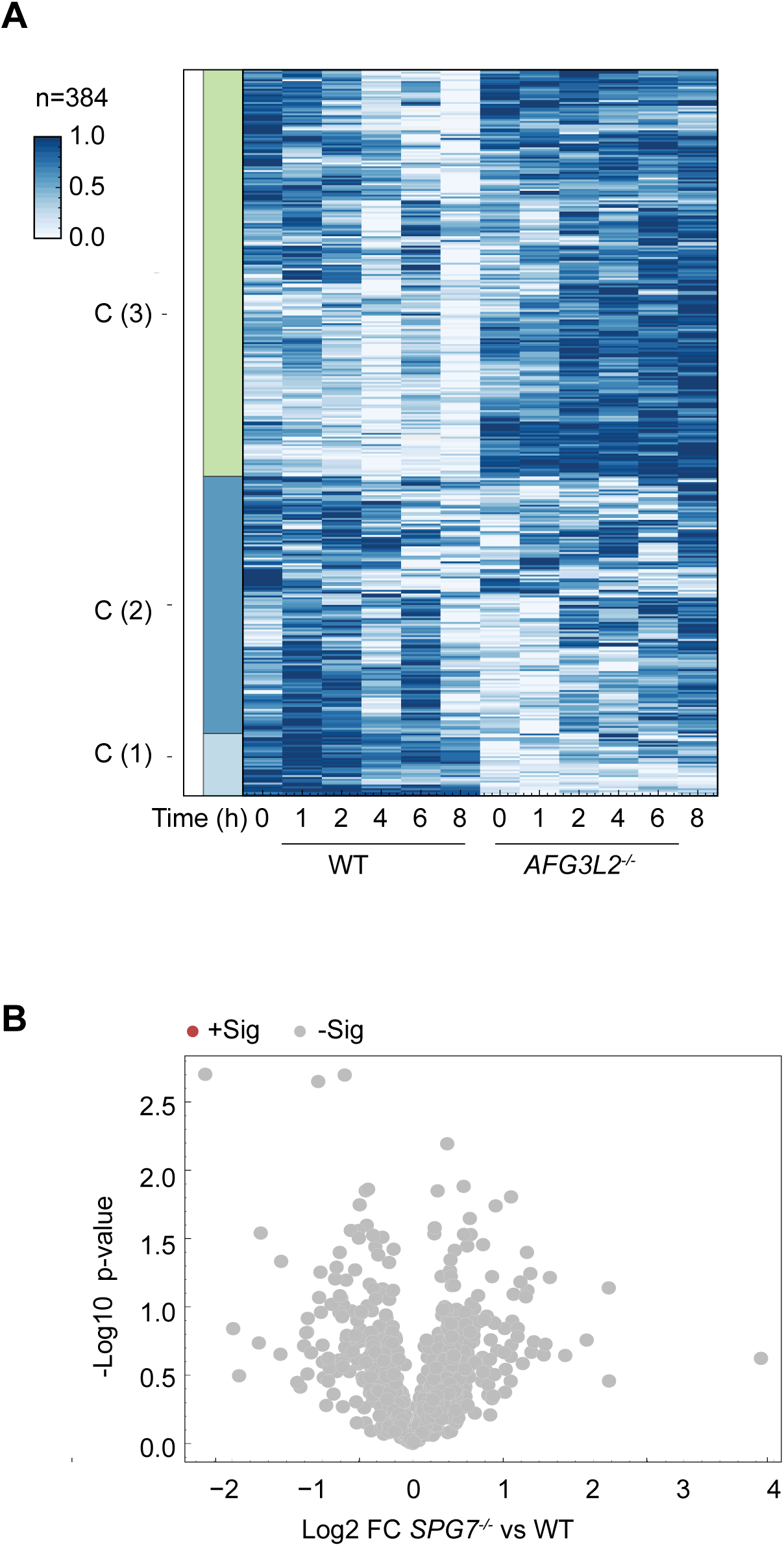
AFG3L2-mediated proteolysis broadly rewires the mitochondrial proteome. **A.** Z-score heatmap showing the log_2_ transformed LFQ intensities scaling the highest intensity to 1 and the lowest intensity to zero after aggregating the biological replicates using the median. Significantly changed proteins (interaction p-value time x genotype <0.01) were identified using a mixed linear model using the time as well as genotype and replicate as variables on the peptide data (random effect). Only proteins with a negative slope (param time) were considered. **B.** Volcano plot depicting the log_2_ fold changes (FC) in protein abundances in *SPG7*^-/-^ versus wildtype (WT) HeLa cells and the negative log_10_ p-value of two-tailed t-tests. n=5 per genotype.

**Supplementary Figure 2.**
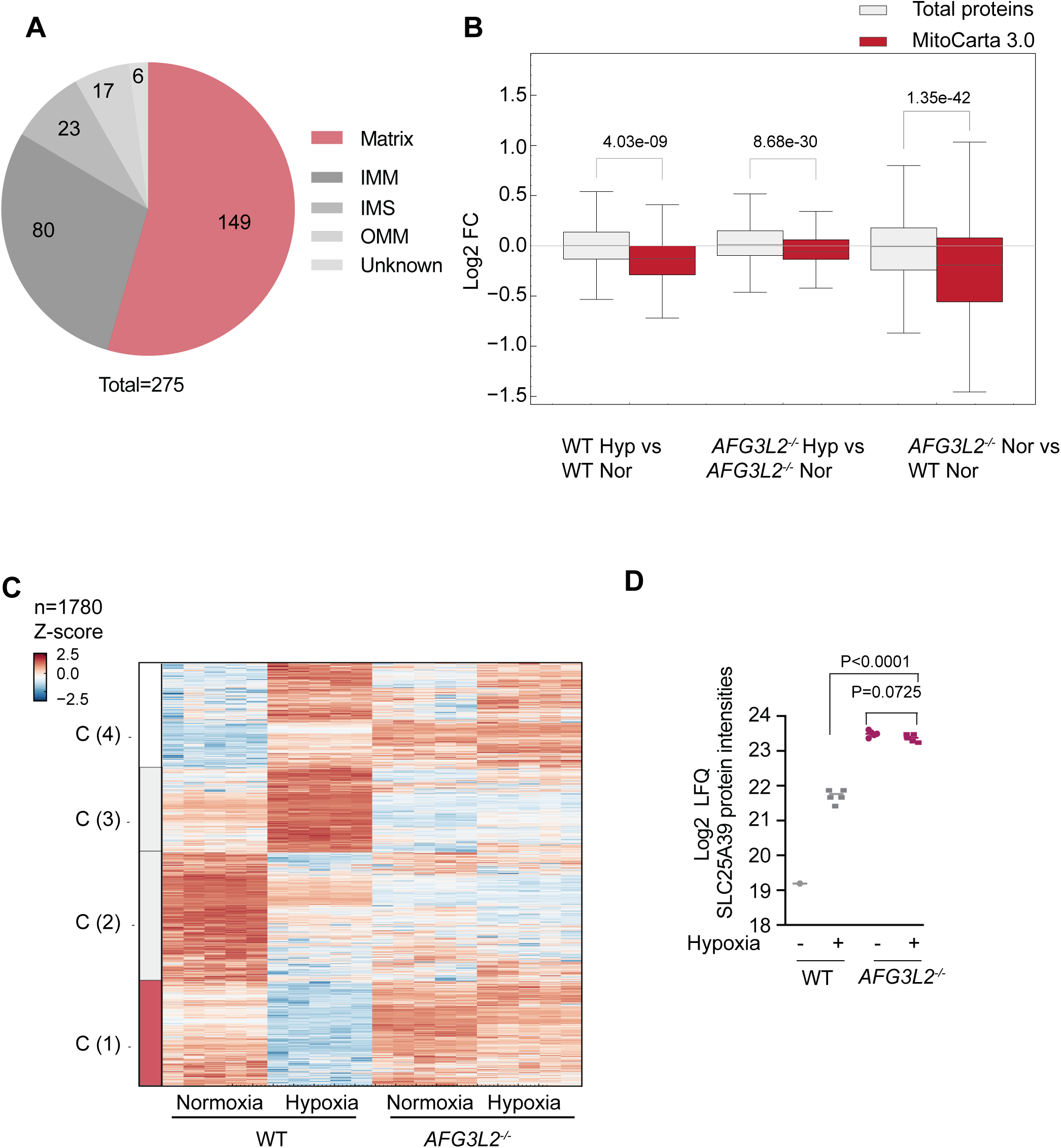
AFG3L2-mediated mitochondrial reprogramming in hypoxia. **A.** Venn-diagram showing the mitochondrial localization of proteins whose abundance is significantly reduced in hypoxia. IMM, inner mitochondrial membrane; IMS, intermembrane space; OMM, outer mitochondrial membrane. **B.** Boxplot analysis of the cellular proteome of wildtype (WT) and *AFG3L2^-/-^*cells in normoxia (Nor) or hypoxia (Hyp). The distribution of relative protein abundances (log_2_ FC) are shown for the total proteome (grey) and for mitochondrial proteins according to MitoCarta3.0 (42). The p-value of a two-tailed t-test is indicated. **C.** Z-score heatmap of log_2_-transformed LFQ intensities of significant proteins in WT and *AFG3L2^-/-^*cells in normoxia and hypoxia (n=5). Indicated proteins were two-way ANOVA significant. **D.** Accumulation of SLC25A39 in *AFG3L2^-/-^* cells.Log_2_-transformed intensities corresponding to SLC25A39 in WT and *AFG3L2^-/-^* cells are shown. The p-values are indicated from the analysis of two-tailed t-test.

**Supplementary Figure 3.**
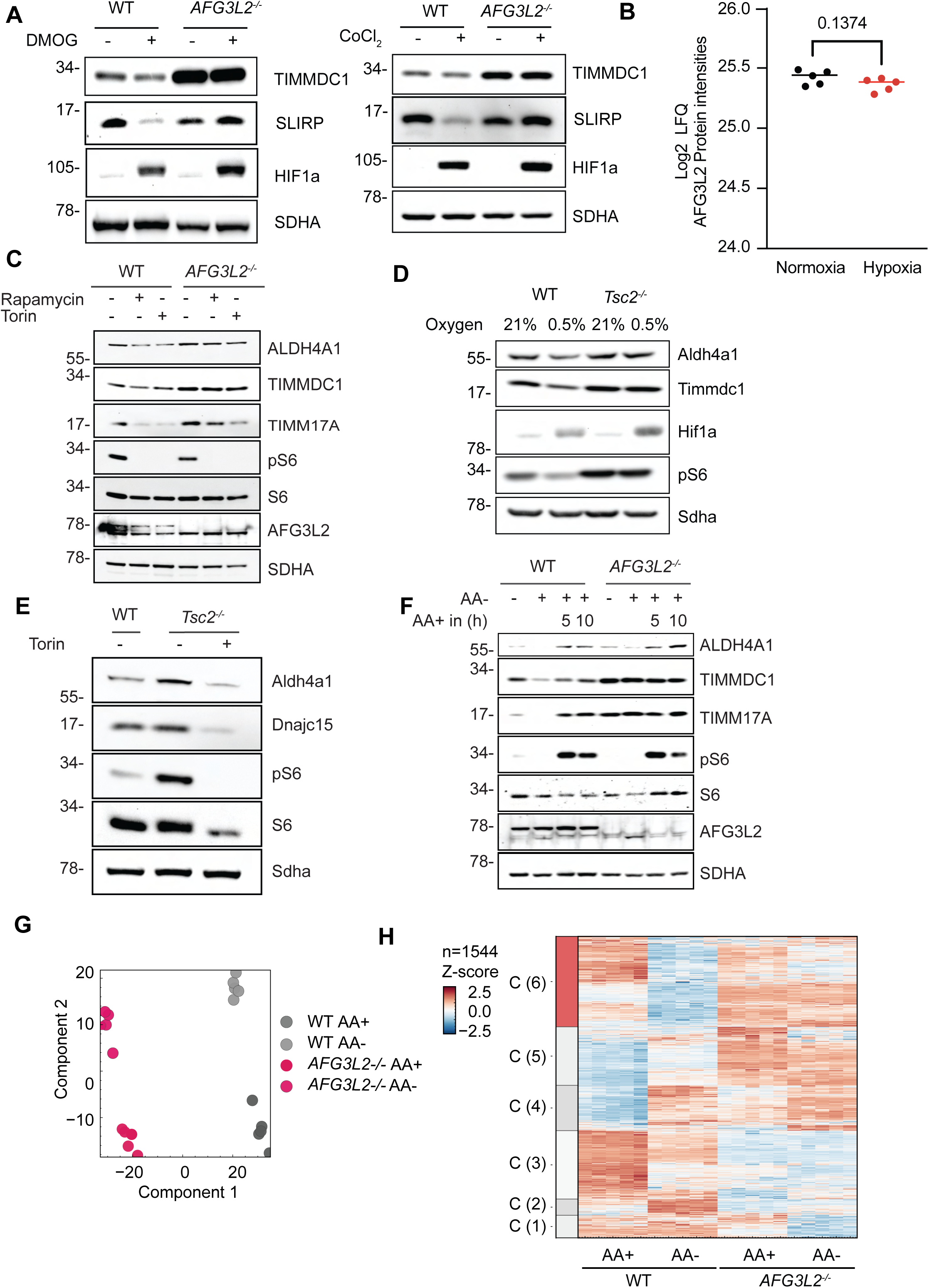
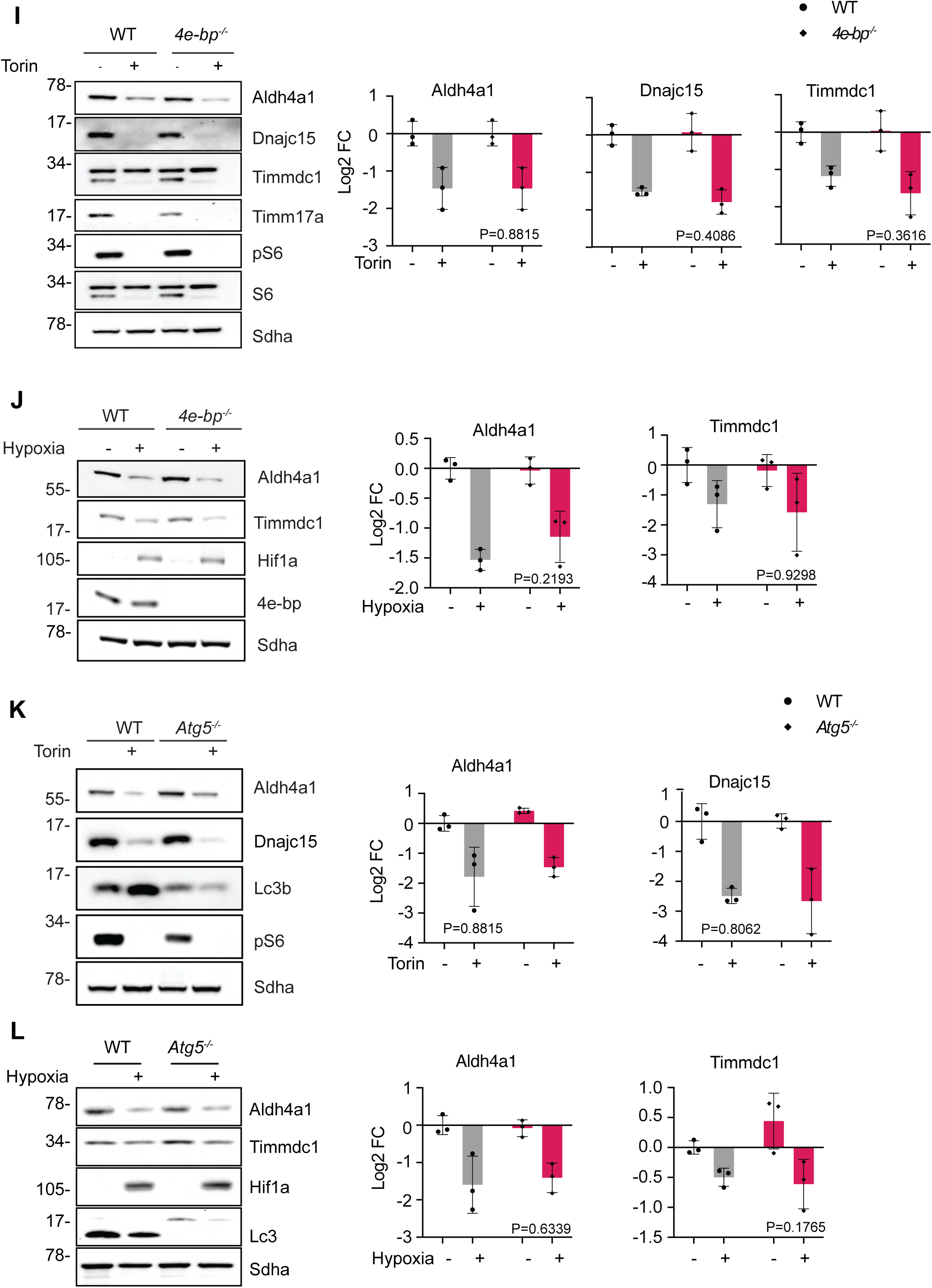
Posttranslational regulation of the AFG3L2-mediated protein turnover. **A.** Immunoblot analysis of WT and *AFG3L2^-/-^* HeLa cells treated with (100 µM) DMOG and (100 µM) CoCl_2_ for 16 hours (n=3). TIMMDC1 and SLIRP are shown as representative AFG3L2 substrates. **B.** Steady-state level of AFG3L2 in normoxia and hypoxia. Log_2_-transformed intensities corresponding to AFG3L2 in normoxia and hypoxia are shown. The p-values are indicated from the analysis of two-tailed t-test. **C.** Immunoblot analysis of WT and *AFG3L2^-/-^* HeLa cells treated with torin (200 nM) or rapamycin (200 nM) for 12 hours (n=3). A representative analysis is shown. **D.** Immunoblot analysis of WT and *Tsc2^-/-^* MEF cells cultered in the presence of indicated oxygen levels (n=3). A representative analysis is shown. **E.** Immunoblot analysis of WT and *Tsc2^-/-^* MEF cells treated with torin (200nM) for 5 hours (n=3). A representative analysis is shown. **F.** Immunoblot analysis of WT and *Tsc2^-/-^* MEF cells treated with torin (200 nM) for 5 hours (n=3). **G.** Principal component analysis (PCA) of proteomic datasets of WT and *AFG3L2^-/-^* cells cultivated in the presence (+AA) or absence (-AA) of amino acids. **H.** Z-score heatmap of log_2_-transformed LFQ intensities of WT and *AFG3L2^-/-^* HeLa cells cultivated in the presence or absence of amino acids (n = 5). (n=1544) are significant in two-way ANOVA analysis. **I.** Immunoblot analysis of WT and *4ebp^-/-^* MEF cells treated with torin (200 nm) for 5 hours. A quantification of steady-state levels of the AFG3L2 substrate proteins Aldh4a1, Dnajc15 and Timmdc1 is shown in the right panels (n=3). The p-value (genotype x treatment) of two-way ANOVA analysis is indicated. **J.** Immunoblot analysis of WT and *4ebp^-/-^* MEF cells cultivated in normoxia and hypoxia for 16hours. A quantification of steady-state levels of the AFG3L2 substrate proteins Aldh4a1 and Timmdc1 is shown in the right panels (n=3). The p-value (genotype x treatment) of two-way ANOVA analysis is indicated. **K.** Immunoblot analysis of WT and *Atg5^-/-^* MEF cells treated with torin (200 nM) for 5 hours. A quantification of steady-state levels of the AFG3L2 substrate proteins Aldh4a1 and Dnajc15 is shown in the right panels (n=3). The p-value (genotype x treatment) of two-way ANOVA analysis is indicated. **L.** Immunoblot analysis of WT and *Atg5^-/-^* MEF cells cultivated in normoxia and hypoxia for 16 hours. A quantification of steady-state levels of the AFG3L2 substrate proteins Aldh4a1 and Timmdc1 is shown in the right panels (n=3). The p-value (genotype x treatment) of two-way ANOVA analysis is indicated.

**Supplementary Figure 4.**
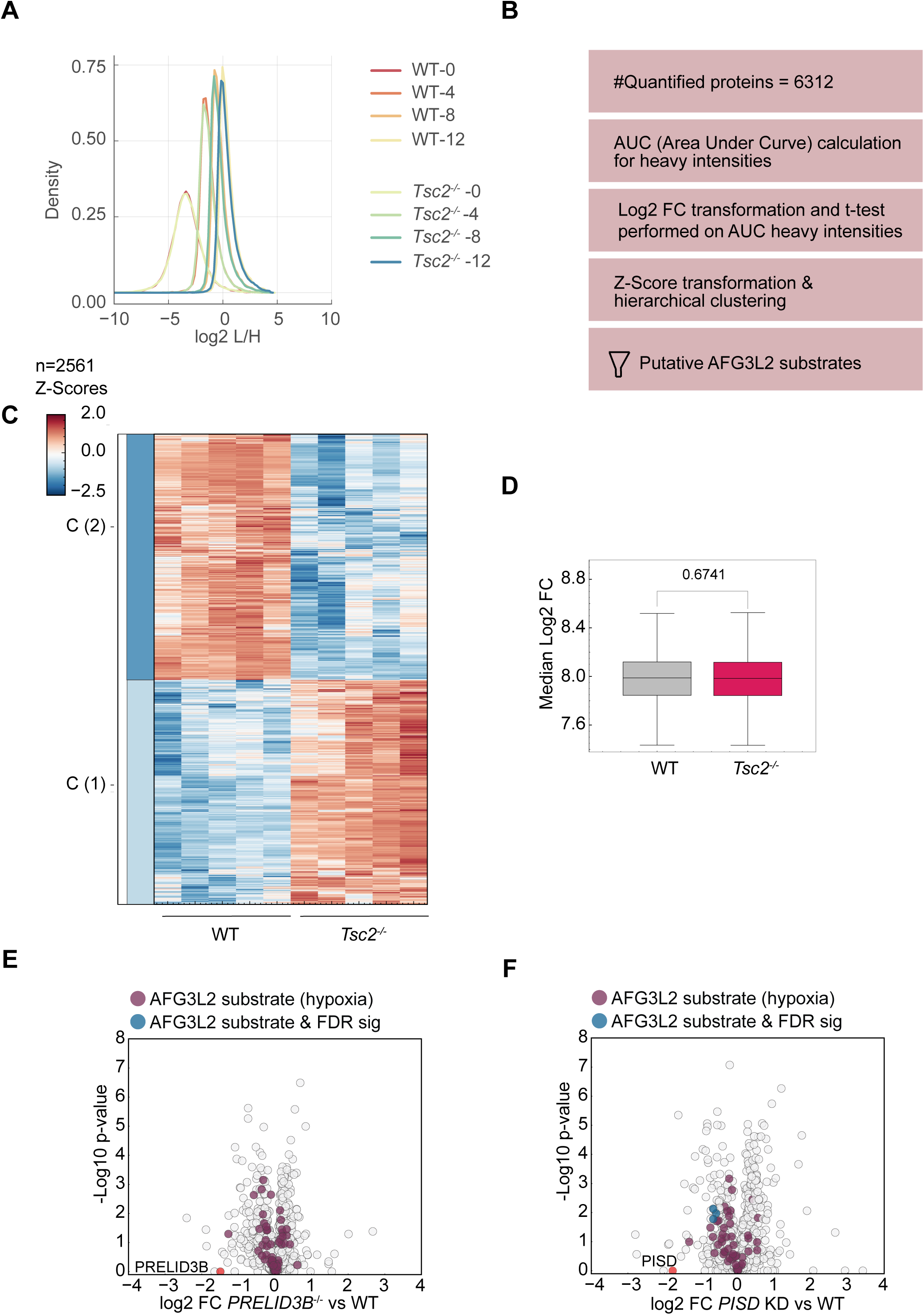
mTORC1 inhibition promotes proteolysis by AFG3L2. **A.** Density plot representing log_2_-transformed ratios of light to heavy intensities of WT and *Tsc2^-/-^* cells at 0, 4, 8 and 12 hours as indicated. **B.** Workflow of the pulsed SILAC proteomic analysis of WT and *Tsc2^-/-^* cells. **C.** Z-score heatmap of log_2_-transformed AUC heavy (N^15^C^13^) intensities of significantly changed proteins in *Tsc2^-/-^* relative to WT cells (n=5). **D.** Boxplot analysis of relative proteins abundances in WT and *Tsc2^-/-^* cells. The distribution of log_2_ FC in the distribution of relative protein abundances (log_2_ FC between *Tsc2^-/-^* and WT cells) are shown for the total proteome. The p-value of a two-tailed t-test is indicated. **E.** Volcano plot depicting the log_2_ fold changes (FC) in protein abundances in *PRELID3B^-/-^* versus WT HeLa cells. AFG3L2 candidate substrate proteins in hypoxia are highlighted in red (n=5). **F.** Volcano plot depicting the log_2_ fold changes (FC) in protein abundances after PISD depletion in WT HeLa cells. AFG3L2 candidate substrate proteins in hypoxia are highlighted in red (n=5).

**Supplementary Figure 5.**
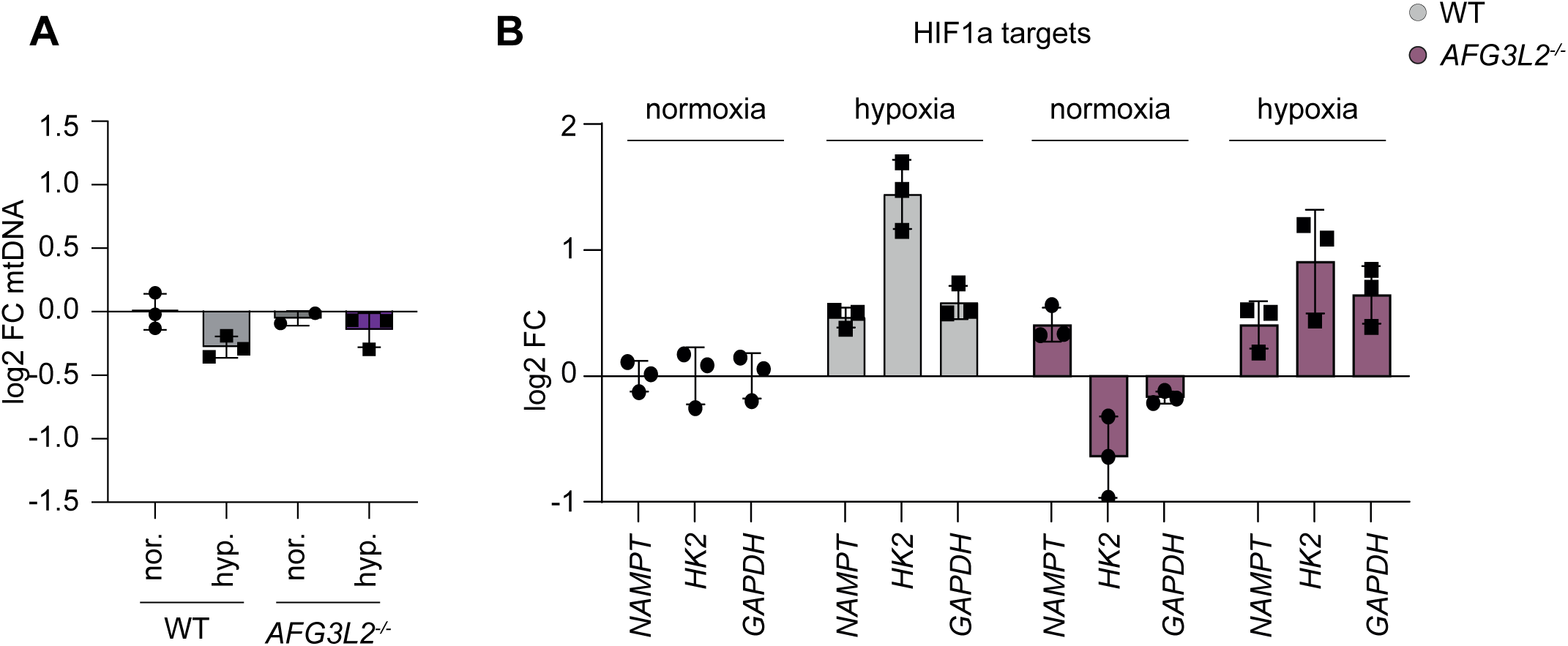
AFG3L2 loss does not affect mtDNA levels nor impair the hypoxic response. **G.** mtDNA level monitored by qPCR of *CYTB* in WT and *AFG3L2^-/-^* cells in normoxia (nor.) or hypoxia (hyp.). Log2 FC relative to WT are shown. (n=3). **H.** Relative mRNA levels of HIF1α target genes analysed by qRT-PCR in WT and *AFG3L2^-/-^* cells in normoxia or hypoxia.

## Supplementary Files

Supplementary File 1. HeLa proteome of AFG3L2 knockout and WT in normoxia/hypoxia including the hierarchical clustering shown in Figure 2D and Suppl. Figure 2C.

**Supplementary File 2.** CHX chase for 1, 2, 4, 6, and 8 hours results in HeLa WT and *AFG3L2*^-/-^ cells

**Supplementary File 3.** Proteomics result of WT and *AFG3L2*^-/-^ w/o amino acids in HeLa cells including the hierarchical clustering shown in Figure 3C and Suppl. Figure 3C.

**Supplementary File 4.** Pulse SILAC proteomics result (heavy intensity over a time course of 4,8 and 12 hours) of WT and *Tsc2*^-/-^ in MEF cells

**Supplementary File 5.** HeLa proteome of *PRELID3B*^-/-^ cells and after PISD knockdown.

